# Understanding the diversity of DNA methylation in Mycobacterium tuberculosis

**DOI:** 10.1101/2020.05.29.117077

**Authors:** Victor Ndhlovu, Anmol Kiran, Derek Sloan, Wilson Mandala, Marriot Nliwasa, Dean B Everett, Mphatso Mwapasa, Konstantina Kontogianni, Mercy Kamdolozi, Elizabeth L Corbett, Maxine Caws, Gerry Davies

## Abstract

Although *Mycobacterium tuberculosis (Mtb)* strains exhibit genomic homology of >99%, there is considerable variation in the phenotype. The underlying mechanisms of phenotypic heterogeneity in *Mtb* are not well understood but epigenetic variation is thought to contribute. At present the methylome of *Mtb* has not been completely characterized. We completed methylomes of 18 *Mycobacterium tuberculosis* (*Mtb*) clinical isolates from Malawi representing the largest number of *Mtb* genomes to be completed in a single study using Single Molecule Real Time (SMRT) sequencing to date. We replicate and confirm four methylation disrupting mutations in lineages of *Mtb*. For the first time we report complete loss of methylation courtesy of C758T (S253L) mutation in the *MamB* gene of Indo-oceanic lineage of *Mtb*. We also conducted a genomic and methylome comparison of the Malawian samples against a global sample. We confirm that methylation in *Mtb* is lineage specific although some unresolved issues still remain.

## Introduction

Tuberculosis (TB) is a disease that remains a global health crisis with an estimated 1.7 billion people infected of which 5-10% will develop the disease in their lifetime (WHO, 2020). The major barriers to disease elimination have been lack of an effective vaccine or fast and effective diagnostic tools, increasing drug resistance and co-infection with HIV (Davies et al., 2014; De Schacht et al., 2019). Mycobacterium tuberculosis (*Mtb*), the causative agent of TB, has a genome with a uniformly high guanine + cytosine (65.6%) owing to minimal incorporation of foreign DNA during its evolution (Cole, 1999). One unique feature of the *Mtb* genome is the large number of genes it contains. Up to 10% of the total coding potential contains polymorphic guanine-cytosine repetitive sequences (PGRS) (Cole, 2002; Grover et al., 2018) which encode two unrelated families of acidic glycine-rich proteins- proline-glutamic acid (PE) and proline-proline glutamic acid (PPE). Specific functions of these genes and their proteins remain unclear (Cole, 2002; Fishbein et al., 2015; J E Phelan et al., 2016) although they have been implicated in immune evasion and virulence (Fishbein et al., 2015; J E Phelan et al., 2016). Consistently, evidence has suggested that proteins located in the cell wall and cell membranes are responsible for diversity in antigenic structure and virulence. This greatly contributes to *Mtb* evolution and adaptation to different hosts (Brennan & Delogu, 2002; Filliol et al., 2006). Although *Mtb* strains have been shown to exhibit genomic homology of >99% (Hershberg et al., 2008) such similarity is rarely replicated in the phenotype. This phenotypic heterogeneity has been seen in the virulence of the *Beijing* strain which has been associated with increasing multidrug resistant TB (MDR-TB) (Cowley et al., 2008; van der Spuy et al., 2009) whereas the East African Indian (EAI) lineage has been associated with lower rates of transmission compared to other lineages (Albanna et al., 2011). Similarly, the Euro-American lineage is the most geographically successful strain (Gagneux & Small, 2007) but specific mechanisms supporting this successful dissemination remain unknown. Phenotypic heterogeneity in *Mtb* has been associated with epigenetic inheritance (Balaban et al., 2004) and the most common epigenetic mechanism in *Mtb* is DNA methylation (Casadesus & Low, 2006; Shell et al., 2013). A few studies have characterized the *Mtb* methylome and revealed three 6-methyladenine (m6A) motifs and their cognate methyltransferases (*Mtases*), *MamA, MamB* and *HsdM* respectively (Shell et al., 2013; Zhu et al., 2015). Using Pacific Biosciences Single Molecule Real Time (SMRT) sequencing, two studies have recently shown that specific mutations in the *Mtases* lead to loss of *Mtase* activity and may play a role in evolution of *Mtb* (J. Phelan et al., 2018; Zhu et al., 2015). At present the methylome of *Mtb* has not been completely characterized, neither has any resulting information been correlated with phenotypic heterogeneity observed in TB patients. Understanding the complete biology of *Mtb* will aid in developing strategies for reducing the *Mtb* treatment duration from the standard 6 months.

We present characterization of methylomes of 18 *Mycobacterium tuberculosis* (*Mtb*) isolates from patients in Blantyre, Malawi including 12 Euro-American lineage (L4) strains, the most prevalent phylogenetic lineage in Malawi, 3 Beijing lineage strains (L2) and 3 Indo-oceanic lineage (L1) strains. This work presents the largest number of *Mtb* genomes of a single lineage to be completed in a single study using Single Molecule Real Time (SMRT) sequencing to date. Additionally, we confirm three confident sequence motifs in *Mtb* and confirm the strain specific mutations responsible for loss of methyltransferase activity in *Mtb*. Additionally, for the first time we report the complete loss of methylation courtesy of a novel mutation C758T (S253L) in Indo-oceanic lineage (L1). Through a genomic and methylome comparative analysis with a global sample of 16 samples we report previously unreported mutation affecting the *pks15/1* locus in L6 and L6 isolates.

## Results

### Lineage Analysis of *Mycobacterium tuberculosis*

Experimental (RD-PCR) and computational (TB-Profiler) outcomes on Malawian strains lineage identification were consistent as: 3/18 (17%) were L1 (Indo-Oceanic), 3/18 (17%) were L2 (East-Asian) and 12/18 (66%) were L4 (Euro-American). *De novo* reporting of global sample lineages (J. Phelan et al., 2018) (16 samples) using TB-Profiler was as follows : 3/16 L1(Indo-oceanic), 2/16 L2 (East-Asian), 3/16 L4 (Euro-American), 2/16 L5 (West African 1 and 6/16 L6 (West African 2) (Table 1). Using a reference with an intact *pks15 (Rv2947c)* gene, it was possible to identify the 15/34 strains belonging to L4 in the combined dataset. These possessed a 7 bp deletion (GGGCCGC) in the *pks15/1* gene as previously documented (Constant et al., 2002; Gagneux & Small, 2007). Additionally, *pks15 (Rv2947c)* could be used to assign lineages to the rest of the samples. All L1 (6/34 strains) had a G1318C substitution and GGGCCGC insertion while L2 (5/34) strains had a GGGCCGC insertion only. All L5 samples had a 9bp deletion (CGGTGCTGG,1097- 1105), a distinct substitution A50G and an insertion GGGCCGC. A L1, L5, L6 (1318 G>C substitution) and a L6 (1658 1bp insertion of G), L1, L2, L5 (1658 7bp insertion) (Fig 1)

**Table 1:** Lineages and sub-lineages of the samples reported by TB-Profiler using assembled genomic sequences (ERS-Malawian and SAMEA-global samples).

| <u>Sample ID</u> | <u>Lineage</u> | <u>Sub-lineage</u> | <u>Sub-sub-lineage</u> |
| --- | --- | --- | --- |
| ERS2711939 | Lineage4 | Lineage4.3 | Lineage4.3.4 |
| ERS2711940 | Lineage4 | Lineage4.3 | Lineage4.3.4 |
| ERS2711941 | Lineage4 | Lineage4.3 |  |
| ERS2711942 | Lineage4 | Lineage4.3 | Lineage4.3.4 |
| ERS2711943 | Lineage1 | Lineage1.1 | Lineage1.1.3 |
| ERS2711944 | Lineage4 | Lineage4.3 | Lineage4.3.4 |
| ERS2711945 | Lineage4 | Lineage4.3 | Lineage4.3.4 |
| ERS2711946 | Lineage4 | Lineage4.3 | Lineage4.3.4 |
| ERS2711947 | Lineage4 | Lineage4.3 | Lineage4.3.4 |
| ERS2711948 | Lineage1 | Lineage1.1 | Lineage1.1.3 |
| ERS2711949 | Lineage4 | Lineage4.3 | Lineage4.3.4 |
| ERS2711950 | Lineage4 | Lineage4.5 |  |
| ERS2711951 | Lineage4 |  | Lineage4.1.2 |
| ERS2711952 | Lineage4 | Lineage4.3 | Lineage4.3.4 |
| ERS2711953 | Lineage2 | Lineage2.2 |  |
| ERS2711954 | Lineage2 | Lineage2.2 |  |
| ERS2711955 | Lineage2 | Lineage2.2 |  |
| ERS2711956 | Lineage1 | Lineage1.1 | Lineage1.1.3 |
| SAMEA104606019 | Lineage1 | Lineage1.1 | Lineage1.1.3 |
| SAMEA104606020 | Lineage1 | Lineage1.1 | Lineage1.1.3 |
| SAMEA104606021 | Lineage1 | Lineage1.1 | Lineage1.1.3 |
| SAMEA104606022 | Lineage5 |  |  |
| SAMEA104606023 | Lineage2 | Lineage2.2 | Lineage2.2.1 |
| SAMEA104606024 | Lineage4 | Lineage4.3 | Lineage4.3.4 |
| SAMEA104606025 | Lineage4 | Lineage4.1 | Lineage4.1.2 |
| SAMEA104606026 | Lineage6 |  |  |
| SAMEA104606027 | Lineage6 |  |  |
| SAMEA104606028 | Lineage2 | Lineage2.2 | Lineage2.2.1 |
| SAMEA104606029 | Lineage4 | Lineage4.3 | Lineage4.3.4 |
| SAMEA104606030 | Lineage5 |  |  |
| SAMEA104606031 | Lineage6 |  |  |
| SAMEA104606032 | Lineage6 |  |  |
| SAMEA104606033 | Lineage6 |  |  |
| SAMEA104606034 | Lineage6 |  |  |

**Figure 1:**
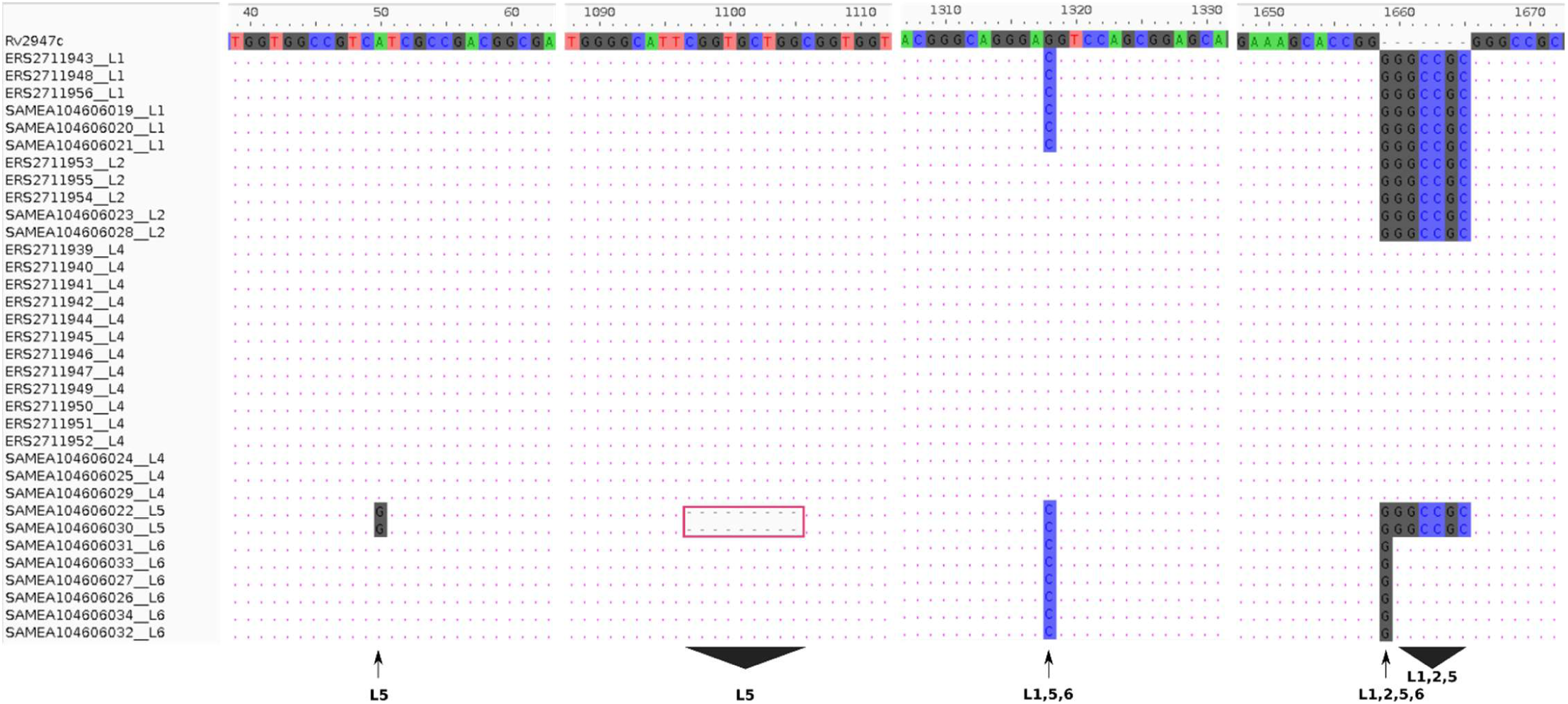
Lineage specific sequence differences relative to the reference gene pks15 (Rv2947c) The pks15 gene from 34 samples was aligned against the reference to display lineage specific variations. Variants were observed in four different locations/ranges within the gene discriminating four lineages L5 (50, A>G substitution), L 5 (1097-1105 CGGTGCTGG deletion), L1, L5, L6 (1318 G>C substitution) and L6 (1658 1 bp insertion of G), L1, L2, L5 (1658 7bp insertion)

### DNA Methylation Patterns

The m6A methylation motifs present in more than 10 isolates were CACGC**<u>A</u>**G (820 sites), C**<u>T</u>**CC**<u>A</u>**G (1947 sites), C**<u>T</u>**GG**<u>A</u>**G (1947 sites), G**<u>A</u>**TN_4_R**<u>T</u>**AC (363 sites) and G**<u>T</u>**AYN_4_**<u>A</u>**TC (363 sites), Motifs C**<u>T</u>**CC**<u>A</u>**G and G**<u>A</u>**TN_4_R**<u>T</u>**AC are paired with C**<u>T</u>**GG**<u>A</u>**G and G**<u>T</u>**AYN_4_**<u>A</u>**TC respectively (S1 Table). No methylation of C**<u>T</u>**GG**<u>A</u>**G and C**<u>T</u>**CC**<u>A</u>**G was reported in all L2 samples including one L6 sample (SAMEA104606027). One sample from L4 (ERS2711941) lacked methylation in C**<u>T</u>**CC**<u>A</u>**G motif (Fig 2A). These motifs are methylated by *mamA Mtase* (Shell et al., 2013). Multiple sequence comparison with the reference gene (*Rv3263*) revealed that all L2 samples had a A809C change resulting in E270A as previously reported (J. Phelan et al., 2018; Shell et al., 2013; Zhu et al., 2015) (including G1199C, W400S in two samples only). Consequently, non-methylated L6 samples had an alteration at A1378G resulting in A460T in *Rv3263*. The L4 sample showing no methylation in motif C**<u>T</u>**CC**<u>A</u>**G had a synonymous substitution at C216T and a non-synonymous substitution at G454A resulting in G152S amino acid substitution. To our knowledge, this potentially methylation disrupting mutation has not been previously reported. The motif CACGC**<u>A</u>**G is methylated by the *mamB Mtase* (J. Phelan et al., 2018; Zhu et al., 2015). Two of the six L1 samples (ERS2711948, ERS2711956) lacked this methylation (Fig 2A). Methylation in the rest of the samples was however below 80% (range 56% – 79.6%) (Fig 2C). Surprisingly, all L1 samples (6) possessed a C758T resulting in amino acid change S253L in the *mamB* gene *(Rv2024c).* This mutation has previously been reported to be responsible for partial loss of methylation in L1 samples (J. Phelan et al., 2018). This is the first time that mutation S253L is being associated with complete loss of *MamB Mtase*. Lineage 2 sample (ERS2711953) and L4 sample (ERS2711945) had low methylation in motif CACGC**<u>A</u>**G (65% and 73 %) compared to other samples from the same lineage (100%) but these samples had no specific mutations in the *mamB* gene. No effect of L6 specific mutation R289C and L5 specific mutation L452V was observed on the mamB methylation in these lineages (Fig 2C). However, non-lineage specific multiple variation was reported at 3’ end. Motifs G**<u>A</u>**TN_4_R**<u>T</u>**AC (363 sites) and G**<u>T</u>**AYN_4_**<u>A</u>** TC (363 sites) are methylated by *hsdM (Rv2756c)* and *hsdS (Rv2761c)* genes (J. Phelan et al., 2018; Zhu et al., 2015). One L1 sample (ERS2711956) showed no methylation in either motif however, ERS2711948 was methylated at G**<u>T</u>**AY N_4_**<u>A</u>**TC only (Fig 2A). The *hsdM* gene sequences were identical for all L1 samples and no 5’-upstream alterations (300bp) were reported either. All the L4 samples lacking G**<u>A</u>**TN_4_RT**<u>A</u>** C/G**<u>T</u>**AYN_4_**<u>A</u>** TC methylation had mutations at T917C resulting in L306P in *hsdM* gene and G74T resulting in G25V amino acid change in *hsdS* gene. While the T917C (L306P) was previously characterized (J. Phelan et al., 2018; Shell et al., 2013; Zhu et al., 2015)., the G74T(G25V) mutation in *hsdS* gene has not been previously reported. The distribution of lineages specific motif methylation is show in Fig 2B and Fig 2D.

**Figure 2:**
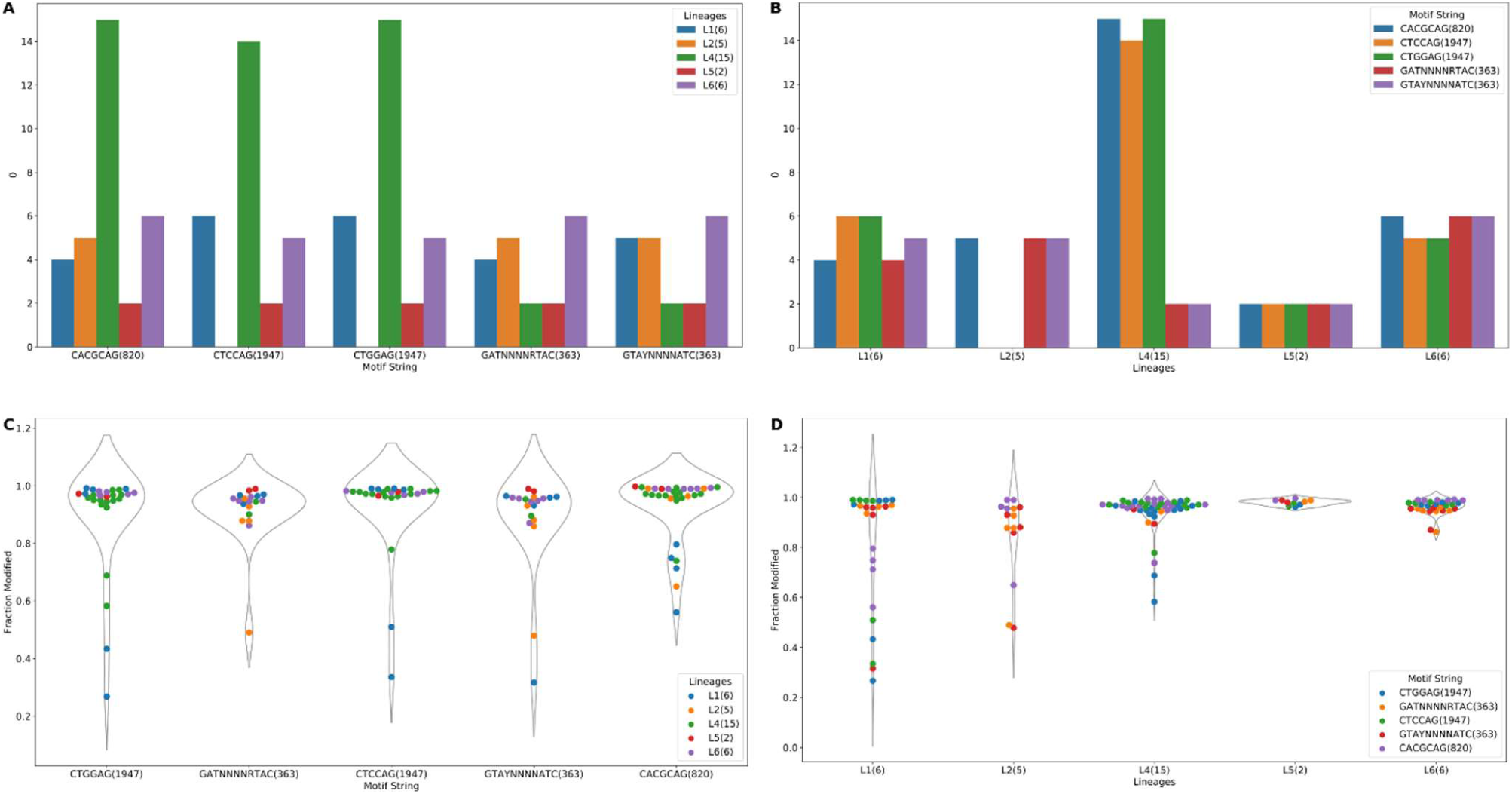
Methylation summary. (A) Distribution of methylated samples in each Lineage for the motifs. (B) Distribution of samples with methylated motifs in each lineage. (C) Methylation efficiency in samples for each motif. (D) Methylation efficiency by motif in each lineage.

### Methylation Efficiency among lineages

Among the samples having methylation in major motifs, most reported methylation efficiency was higher than 82% (Fig 2C, S1 Table). Lineage 4 sample (ERS2711941) had 58% methylation for C**<u>T</u>**CC**<u>A</u>**G and it lacked methylation on the C**<u>T</u>**GG**<u>A</u>**G motif while another L4 sample (ERS2711945) had 69% and 79% methylation on C**<u>T</u>**GG**<u>A</u>**G and C**<u>T</u>**CC**<u>A</u>**G respectively. Two L1 samples (ERS2711948, ERS2711956) showing methylation of 27% and 36%, 43% and 51% for C**<u>T</u>**GG**<u>A</u>**G and C**<u>T</u>**CC**<u>A</u>**G respectively but having no specific mutation in methylation conferring genes. Methylation distribution within motifs for each sample is displayed in Fig 2D. Lineage 1 sample (ERS2711948) was methylated at 32% on motif G**<u>T</u>**AYN_4_**<u>A</u>**TC, while L2 sample (ERS2711953) was methylated at 48% on this motif and 49% on motif G**<u>A</u>**TN_4_R**<u>T</u>**AC. Other samples with low efficiency were as follows: ERS2711953 from L2 with 65% methylation efficiency and L1 samples SAMEA104606020, SAMEA104606019, SAMEA104606021 and ERS2711943 with 71%, 80%, 75% and 56% respectively on CACGCAG (Fig 2C and Fig 2D).

### Comparison of Methylation within *Mtb* strains

The strain arrangements in the m6A IPD ratio based cladogram clusters and genome based maximum likelihood (ML) phylogeny were compared (Fig 3). The samples clustered into four IPD based groups. However, in the ML phylogeny lineages formed distinct clusters. Lineage 2 samples and one L6 sample (SAMEA104606027) with no C**<u>T</u>**GG**<u>A</u>**G and C**<u>T</u>**CC**<u>A</u>**G methylation clustered together. Two samples belonging to L4 (ERS2711951, SAMEA104606025) having no methylation in G**<u>A</u>**TN_4_R**<u>T</u>**AC and G**<u>T</u>**AYN_4_**<u>A</u>**TC motifs clustered with L5 and L6 samples in the IPD ratio cladogram (Fig 3). In the recombination hotspot check, 256 genes were reported to have been affected (Fig 4). Ninety-one well annotated genes were affected due to insertions/deletions (Indels) varying size from 1 to 36016pb affecting a large number of PPE family (21), PE-PGRS family (25) and ESAT 6 (6) genes. Lineage specific recombination relative rate to mutation ratio (r/m) reported as L1: 0.968310, L2:1.865780, L4:4.915385, L5:1.001656, L6:1.066062.

**Figure 3.**
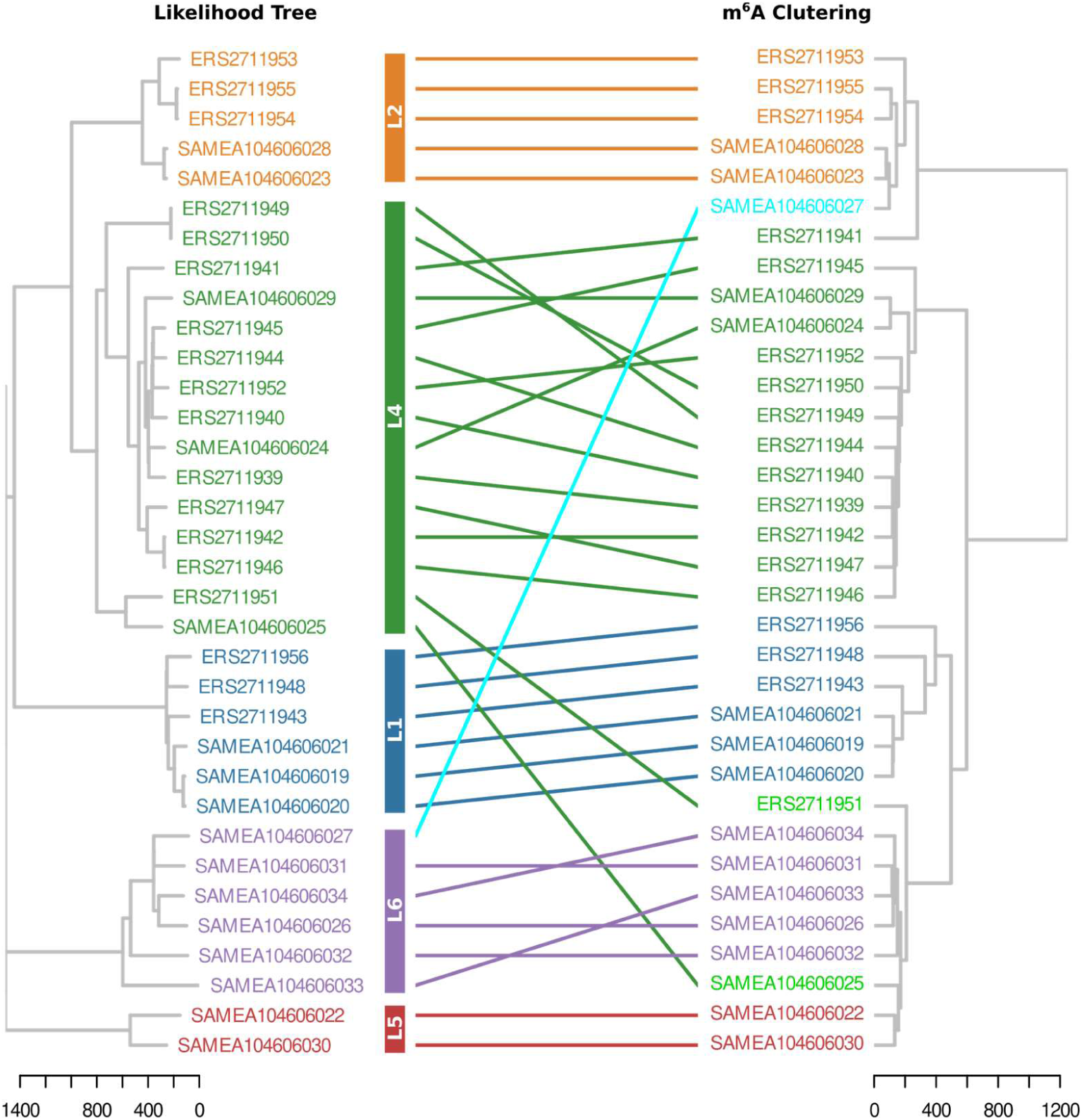
Tanglegram of hierarchically clustered samples. Clustering was based on IPD and ML phylogeny. Samples are coloured based on lineages. Three samples clustered separately from their lineage.

**Figure 4.**
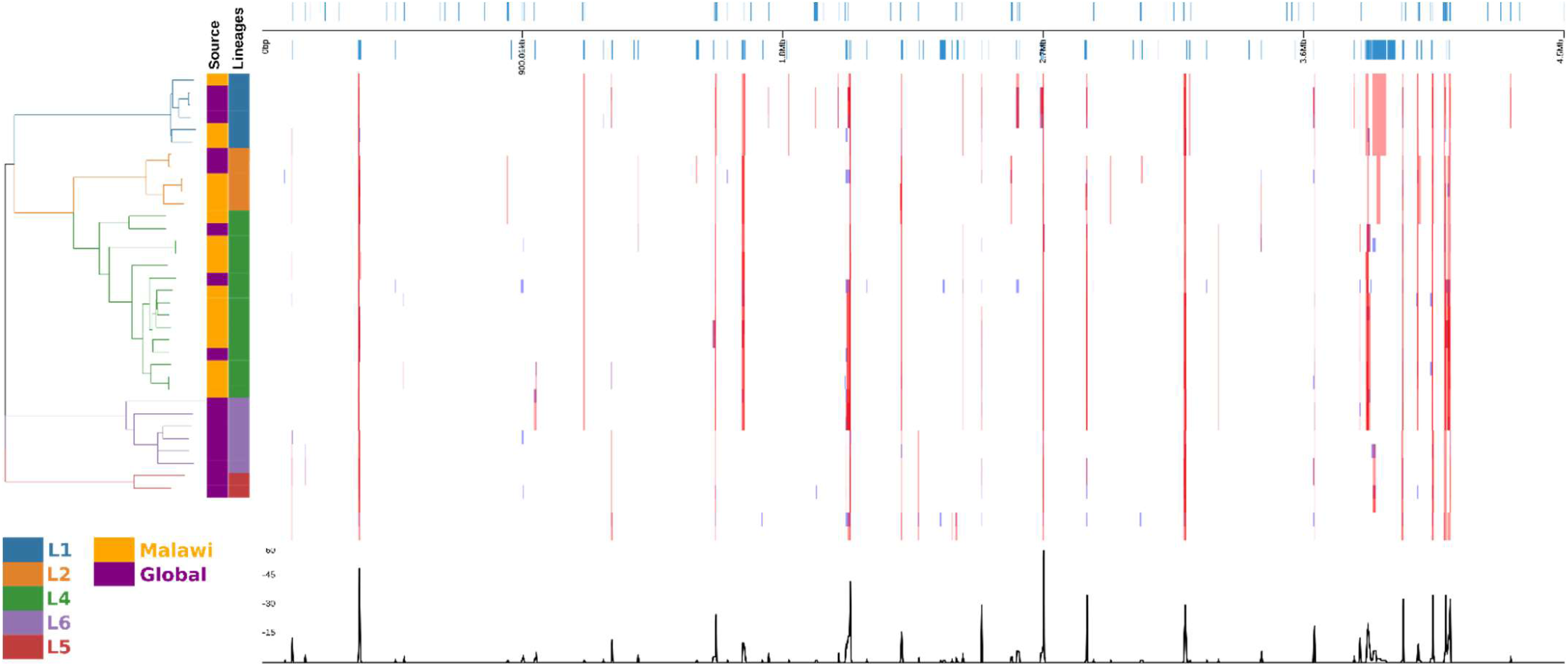
Diverse region in different samples and lineages. Differences are displayed in alignment frame of the different samples and lineages calculated with default Gubbins parameters. Regions of affected gene locations in the alignment (top). The phylogeny of the 34 samples (left). Recombination events (bottom)

### Stability of methylation within Mycobacterium tuberculosis strains

It was important to study the effect of culture media on methylation patterns. Among our sequences isolates, two (ERS2711943 and ERS2711952) were MGIT grown, the methylation pattern did not appear distinct from the same lineages except “CACGC**<u>A</u>**G” for ERS2711943 was the lowest in the L1 samples, G**<u>A</u>**TN_4_R**<u>T</u>**AC was detected in ERS2711943 only. No significant difference could be established between liquid and solid culture isolates for methylated motif C**<u>T</u>**GG**<u>A</u>**G (Fishers exact test p=0.76). As for motif CACGC**<u>A</u>**G solid cultured isolates were methylated at an average 76% while liquid cultures were methylated an average 97%. It was found that liquid cultured isolates were significantly more methylated than solid cultures (Fisher’s exact p=0.02) for motif CACGC**<u>A</u>**G. It was observed that this difference was as a result of sample ERS2711943 being lowly methylated at 56% compared to the rest at >95%.

We next investigated methylation within the gene regions and promoter regions of genes. Methylation within gene regions ranged between 49% to 51% in each strand and there was no over representation of methylation by strand (Chi squared test with Yates correction P=0.44). On the other hand, methylation within promoter regions of genes ranged from 37% to 62% by strand of the promoter methylation. Again there was no significant statistical differences observed by strand (Fishers exact test P=0.19).

## Discussion

We sequenced 18 genomes of clinical *Mtb* isolates from Blantyre, Malawi using SMRT sequencing technology and analysed them along with a set of 16 global samples. Studies of *Mtb* DNA methylation using SMRT sequencing have focused on strains originating from the United States of America (Shell et al., 2013; Zhu et al., 2015), Asia (Zhu et al., 2015) and more recently a small global sample that included Europe, Asia, West Africa and South Africa (J. Phelan et al., 2018). To date no *Mtb* samples from Malawi or the surrounding region have been subjected to either PacBio SMRT sequencing technology or DNA methylation analysis. In our study, SMRT sequencing of 18 *Mtb* clinical isolates from Malawi revealed three confidently identified *Mtase* across the three lineages under study. The activity of these *Mtase* could be inactivated by three different mutations somewhat in a lineage specific manner. The *Mtase MamA* was found to be active in all isolates except three L2 (*Beijing)* isolates putatively courtesy of a point mutation A809C (E270A). This point mutation has been previously characterized (Shell et al., 2013).

Interestingly, L2 (*Beijing)* strains have a higher propensity to cause active disease and have been associated with increasing drug resistance in some geographical areas (Cowley et al., 2008; van der Spuy et al., 2009). Whether loss of this *Mtase* could be associated with success of this organism is an area of interest for future studies. A recent study however failed to establish a possible role of methylation in virulence of *Beijing* strains (*Computational characterisation of DNA methylomes in mycobacterium tuberculosis Beijing hyper- and hypo-virulent strains*, n.d.). Similarly, the *MamB Mtase* (motif CACGC**<u>A</u>**G) was absent in two (L1) Indo-oceanic isolates. This could be attributed to a C758T (S253L) novel missense mutation recently characterized elsewhere (J. Phelan et al., 2018) and confirmed in this study. While this mutation was putatively found to lead to partial methylation (50-60%) in a previous study, for the first time, we report that it could also lead to complete loss of *Mtase* activity as two of our L1 isolates had 0% methylation. And whether indeed this mutation is responsible for this partial/total loss of methylation now remains debatable. This mutation is present only in EAI6 family of L1 which have been shown to be responsible for recent TB outbreaks globally (Duarte et al., 2017). It is still unknown whether the C758T (S253L) mutation contributes to this transmission. Our investigations as to how the mutation C758T (S253L) could lead to partial loss of methylation in one sample and complete loss in others yielded nothing as we found the rest of the *mamB gene* to be identical in all the L1 samples including the global samples. There could be yet other unknown mechanisms, possibly a second gene regulating this methylation. In L4 isolates lack of *HsdM* methylation could be attributed to the C917T (P306L) mutation which was present in 11/12 Malawian isolates. Again lack of methylation was associated with this mutation in all L4 global samples. These results are consistent with previous studies which seem to suggest that the P306L mutation is very common in L4 strains (Shell et al., 2013; Zhu et al., 2015). In one study, the mutation was found to be present in 35 out of 37 isolates L4 clinical isolates (Zhu et al., 2015). No cognate restriction enzyme for *HsdM* has been identified suggesting it could be an orphan *Mtase* (Zhu et al., 2015). Its principal function could therefore be related to gene regulation rather than restriction modification. Lineage 4 isolates have the highest global prevalence than any other lineage and more studies will be required to establish whether loss of *HsdM* methylation could be associated with this global success. If indeed *HsdM Mtase* is disrupted by this mutation in L4, it remains intriguing how some L1 isolates could lose *HsdM Mtase* in absence of P306L mutation or any other mutation in the *hsdM* gene. The high frequency of *Mtase* disrupting mutations in *Mtb* could be suggestive of a competitive fitness advantage such as immune evasion or even persistence. We found the efficiency of *Mtases* to be highly variable within and across lineages even in presence of a *Mtase* gene. The polyketide synthase (*pks15/1*) locus is responsible for biosynthesis of phenolic glycolipid (PGL), a cell wall component (Caws et al., 2008; Reed et al., 2004) and has widely been used to discriminate between L4 isolates against L1 and L2 isolates owing to a 7bp deletion in L4 isolates (Caws et al., 2008; Gagneux & Small, 2007). In this study for the first time, we have demonstrated the potential of using the *pks15/1* locus to classify L5 isolates using 9bp (CGGTGCTGG) deletion a distinct substitution A50G and an insertion GGGCCGC while L6 isolates could also be classified using a 6bp (GGGCCGC) at the same position of the 7bp deletion in L4 isolates. The *pks15/1* locus therefore could be a valuable marker for identifying isolates belonging to L5 and L6. The large number of genomic re-arrangements observed in mostly cell wall component genes PPE, PE-PGRS and ESAT-6 is evidence of the large variations that exist among different strains and lineages of *Mtb* in responding to host immunity.

We believe the complete characterization of DNA methylation in *Mtb* could help provide clues to some of the clinical phenotypes which have been associated with strain and lineage variation. In this study no compelling correlation could be established between methylation and *Mtb* growth condition although MGIT cultures were shown to sequence at a slightly lower coverage. Overall data presented in this study shows the potential of SMRT sequencing long reads to help us better understand the complete biology of *Mycobacterium tuberculosis* by resolving difficult regions of the genome and elucidating the complete methylome of the pathogen. This study could not establish the direct association between mutations and loss of *Mtase* activity and also why some samples could show low levels of *Mtase* activity than others. To better understand the complete impact of DNA methylation within specific strains and lineages, subsequent studies will need to integrate transcriptomic and proteomic data to methylomes.

## Materials and methods

### Sample collection

Frozen archived clinical isolates from a previous prospective cohort study, Studying Persistence and Understanding Tuberculosis in Malawi (S.P.U.T.U.M) (Sloan et al., 2015) were characterized. These were from patients aged 16-65 years old presenting with bacteriologically culture confirmed pulmonary *Mtb* between June 2010 and December 2011 at Queen Elizabeth Central Hospital in Blantyre, Malawi. Out of a total of 133 *Mtb* positive isolates, 18 were selected based on which isolates were the first to be successfully revived from frozen state and used in this study.

### Bacterial growth conditions

All experiments involving *Mtb* were performed in a Biosafety Level (BSL) 3 Laboratory, University of Malawi-College of Medicine/ Malawi Liverpool Welcome Trust (CoM/MLW) TB laboratory and at Liverpool School of Tropical Medicine following Standard Operating Procedures (SOPs). All reagents used were from Sigma-Aldrich unless otherwise stated. For liquid culture, strains were grown in Middlebrook 7H9 broth base supplemented with oleic acid, albumin, dextrose and catalase (OADC) and an antibiotic mixture of polymyxin B, amphotericin B, nalidixic acid, trimethoprim and azlocillin (PANTA). Tubes were incubated in a BACTEC MGIT 960 instrument at 37°C and monitored once a week for possible growth for up to eight weeks. Isolates used in the study were from a previous study for which ethics approval had previously been granted by the College of Medicine Ethics Committee (COMREC), University of Malawi (Sloan et al., 2015). Solid culture inoculation was done on Lowenstein-Jensen (LJ) slopes following laboratory SOP. Cultures were grown to mid-log phase and harvested at ∼7th week and used for DNA isolation. *Mtb* was confirmed using both the BD MGIT TBC ID test device (Becton Dickinson, Maryland U.S.A) following manufacturer’s instructions and Ziehl Neelsen (ZN) staining for acid fast bacilli (AFB).

### DNA Extraction

Genomic DNA was isolated using the traditional Cetyltrimethylammonium bromide (CTAB) method as previously described (Somerville et al., 2005). Extracted DNA was quantified using Qubit 3.0 fluorometer (Life Technologies, USA) according to manufacturer’s instructions and DNA purity was determined on a NanoDrop ND-1000 Spectrophotometer V3.7 (Thermo Scientific, Wilmington U.S.A) following manufacturer’s instructions. DNA purity was checked at absorbance 260nm and 280nm by calculating a ratio of A260/A280. DNA quality was analyzed on 1.5% Agarose Gel electrophoresis and visualized under UV light following ethidium bromide staining.

### Genotyping of *Mtb* Isolates

Genotyping of isolates was done at the Liverpool School of Tropical Medicine, United Kingdom. Lineage specific deletions were detected using a singleplex PCR based method with specific oligonucleotide primers targeting the regions of difference RD239, RD105 and RD750. PCR reactions were performed as documented in our previous publication (Ndhlovu et al., 2019).

### DNA Sequencing

Purified genomic DNA libraries were sequenced at the Centre for Genomic Research (CGR), Institute of Integrative Biology, University of Liverpool, United Kingdom. DNA libraries were purified with 1x cleaned AMPure beads (Agencourt) and the quantity and quality was assessed using the Qubit and NanoDrop assays respectively. In addition, the Fragment Analyzer using a high sensitivity genomic kit (Advanced Analytical Technologies, Inc.) was used to determine the average size of the DNA and the extent of degradation. DNA was treated with Exonuclease V11 at 37 **°**C for 15 minutes. The ends of the DNA were repaired as described by the manufacturer (Pacific Biosciences, Menlo Park, CA, USA). The sample was incubated for 20 minutes at 37 **°**C with DNA damage repair mix supplied in the SMRTbell library kit (Pac Bio). This was followed by a 5-minute incubation at 25 **°**C with end repair mix. DNA was cleaned using 0.5x AMPure and 70% ethanol washes. DNA was ligated to adapter overnight at 25 **°**C. Ligation was terminated by incubation at 65**°**C for 10 minutes followed by exonuclease treatment for 1 hour at 37**°**C. The SMRTbell library was purified with 0.5x AMPure beads. The library was size selected with 0.75% blue pippin cassettes in the range 7000-20000 bp. The recovered fragments were damage repaired again. The quantity of library and therefore the recovery was determined by Qubit assay and the average fragment size determined by Fragment Analyzer. SMRTbell library was annealed to sequencing primer at values predetermined by the Binding Calculator (PacBio) and a complex made with the DNA polymerase (P6/C4 chemistry). The complex was bound to Magbeads and this was used to set up the required number of SMRT cells for the project (two for each sample). Sequencing was performed on Pacific Biosciences RSII sequencing system (Pacific Biosciences, Menlo Park, CA, USA) using 360-minute movie times per cell, yielding ∼ 300x average genome coverage. The generated data have been submitted to the ENA databases (Bio-Project: <u>PRJEB28592</u>).

### Bioinformatics Analysis

Generated long Pacbio reads were analysed using the RS_Modification_and_Motif_Analysis.1 protocol as part of SMRT analysis in SMRT Portal (version 2.2.0). To increase the robustness of our analysis, we included previously published *Mtb* methylation study Pacbio data (Bio-project: PRJEB21888) (J. Phelan et al., 2018) and conducted both genomic and methylation comparisons of the two datasets. Although Bio-project PRJEB21888 had 18 genomes, we could only access 16 and these were used in our analysis. However, we evaluated PRJEB21888 sequences using SMRT Portal (version 5.1.0). Reads were mapped using the Basic Local Alignment with Successive Refinement (BLASR) (Chaisson & Tesler, 2012) algorithm within the SMRT portal. Strain specific genomes were generated by mapping the reads to the reference genome (H37Rv) using Quiver tool. Standard settings were used to detect base modifications and methylation motifs in the strain’s genome. Inter-pulse duration (IPD) ratio (observed vs expected) was measured for the modification detection (Zhu et al., 2015). Computational validation of our samples’ lineages and lineage identification of PRJEB21888 samples were done using TB-Profiler (Jody E Phelan et al., n.d.). Comparative analysis of *pks15 (Rv2947c)* gene was used to report lineages of the samples specifically those from PRJEB21888. The MAFFT (version 7.310) (Katoh & Standley, 2013) was used to generate multiple sequence alignment of consensus sequences against H37Rv reference. Following removal of the reference from the alignment, maximum likelihood (ML) phylogeny was constructed for the remaining sequences using RaxML (v8.2.12) GTR+Γ model (Stamatakis, 2014) applying 1000 bootstrap iterations. Although *Mtb* has a highly rigid and non-recombinogenic genome (>99% nucleotide identity), to report diverse genomic regions among isolates, Gubbins (2.4.1) (Croucher et al., 2015) was applied with the default parameters over previously generated alignment of 34 genomes and earlier constructed ML phylogeny as an initial tree. Identified recombination hot spots were plotted with phylogeny generated without hot spots, affected genes details and the metadata using Phandango (Hadfield et al., 2018). Samples were clustered hierarchically based on m6A IPD ratio pattern.

Multiple sequence alignment of *Mtase* genes (*mamA, mamB, hsdM* and *hsdS*) sequences against the reference gene from H37Rv genome was used to identify possible mutations responsible for loss of methylation. Comparative analysis of well characterized methylation sites among samples were performed. Clustering of the samples based on their reported IPD ratios at methylated sites was performed and compared with clustering in ML phylogeny.

## Acknowledgements

We thank the guardians and patients who participated in this study, and the staff at Queen Elizabeth Central Hospital for their assistance.

## Supplementary files

Supplementary Table 1. Methylation efficiency for 34 Mycobacterium tuberculosis samples

