## Supporting Table 1 for "Understanding the diversity of DNA methylation in Mycobacterium tuberculosis"

**Supporting Table 1: Methylation efficiency for 34 Mycobacterium tuberculosis samples**

| Sample | Lineage | CACGCAG 820) | CTCCAG (1947) | CTGGAG (1947) | GATNNNNRTAC (363) | GTAYNNNNATC (363) |
|---|---|---|---|---|---|---|
| ERS2711939 | L4 | 0.96585363 | 0.972265 | 0.94966614 | 0 | 0 |
| ERS2711940 | L4 | 0.9792683 | 0.97842836 | 0.9686698 | 0 | 0 |
| ERS2711941 | L4 | 0.96707314 | 0 | 0.58294815 | 0 | 0 |
| ERS2711942 | L4 | 0.9768293 | 0.98151004 | 0.96712893 | 0 | 0 |
| ERS2711943 | L1 | 0.5609756 | 0.9866461 | 0.972265 | 0.93663913 | 0.93112946 |
| ERS2711944 | L4 | 0.96585363 | 0.9712378 | 0.94453007 | 0 | 0 |
| ERS2711945 | L4 | 0.7390244 | 0.7786338 | 0.68875194 | 0 | 0 |
| ERS2711946 | L4 | 0.96829265 | 0.97740114 | 0.95428866 | 0 | 0 |
| ERS2711947 | L4 | 0.97195125 | 0.9712378 | 0.9481253 | 0 | 0 |
| ERS2711948 | L1 | 0 | 0.5100154 | 0.43348742 | 0 | 0.3168044 |
| ERS2711949 | L4 | 0.95731705 | 0.9583975 | 0.9244992 | 0 | 0 |
| ERS2711950 | L4 | 0.97195125 | 0.9820236 | 0.9583975 | 0 | 0 |
| ERS2711951 | L4 | 0.9646341 | 0.9794556 | 0.95017976 | 0.90082645 | 0.8953168 |
| ERS2711952 | L4 | 0.9487805 | 0.96302 | 0.9342578 | 0 | 0 |
| ERS2711953 | L2 | 0.65 | 0 | 0 | 0.4903581 | 0.47933885 |
| ERS2711954 | L2 | 0.95609754 | 0 | 0 | 0.8787879 | 0.8595041 |
| ERS2711955 | L2 | 0.9634146 | 0 | 0 | 0.8787879 | 0.8815427 |
| ERS2711956 | L1 | 0 | 0.33538777 | 0.26759118 | 0 | 0 |
| SAMEA104606019 | L1 | 0.7963415 | 0.990755 | 0.99126863 | 0.969697 | 0.9641873 |
| SAMEA104606020 | L1 | 0.7134146 | 0.9902414 | 0.9892142 | 0.96694213 | 0.9614325 |
| SAMEA104606021 | L1 | 0.7487805 | 0.98767334 | 0.9871597 | 0.9641873 | 0.9586777 |
| SAMEA104606022 | L5 | 0.997561 | 0.97842836 | 0.972265 | 0.9889807 | 0.9889807 |
| SAMEA104606023 | L2 | 0.9902439 | 0 | 0 | 0.95592284 | 0.9614325 |
| SAMEA104606024 | L4 | 0.99512196 | 0.98870057 | 0.9851053 | 0 | 0 |
| SAMEA104606025 | L4 | 0.99268293 | 0.96302 | 0.96764255 | 0.94490355 | 0.95316803 |
| SAMEA104606026 | L6 | 0.9902439 | 0.978942 | 0.97791475 | 0.9476584 | 0.95592284 |
| SAMEA104606027 | L6 | 0.9890244 | 0 | 0 | 0.862259 | 0.8705234 |
| SAMEA104606028 | L2 | 0.9902439 | 0 | 0 | 0.92837465 | 0.93112946 |

**Supporting Table 1: Methylation efficiency for 34 Mycobacterium tuberculosis samples**

| | | | | | | |
|---|---|---|---|---|---|---|
| SAMEA104606029 | L4 | 0.99512196 | 0.9892142 | 0.9861325 | 0 | 0 |
| SAMEA104606030 | L5 | 0.9890244 | 0.9650745 | 0.9619928 | 0.9834711 | 0.9807162 |
| SAMEA104606031 | L6 | 0.9890244 | 0.9820236 | 0.98099643 | 0.9614325 | 0.95592284 |
| SAMEA104606032 | L6 | 0.9890244 | 0.972265 | 0.97072417 | 0.94490355 | 0.9476584 |
| SAMEA104606033 | L6 | 0.99268293 | 0.9763739 | 0.9753467 | 0.95592284 | 0.9586777 |
| SAMEA104606034 | L6 | 0.9902439 | 0.96815616 | 0.9661017 | 0.9476584 | 0.94490355 |
